# An odor delivery system for arbitrary time-varying patterns of odors, mixtures and concentrations

**DOI:** 10.1101/077875

**Authors:** Priyanka Gupta, Dinu F Albeanu, Upinder S Bhalla

**Affiliations:** National Center for Biological Sciences, Bangalore, Karnataka 560065, India; Cold Spring Harbor Laboratory, Cold Spring Harbor, NY 11724, USA; Watson School of Biological Sciences, Cold Spring Harbor, NY 11724, USA

## Abstract

Odor stimuli in the natural environment are intermittent and the concentration of any given odor fluctuates rapidly over time. Further, even in the simplest scenario, the olfactory sensors receive uncorrelated, intermittent inputs in the form of odor plumes arising from several odor sources in the local environment. However, typically used odor stimuli under laboratory settings are restricted to long-duration (~seconds), single pulse of one odor at a time that are rarely encountered in nature. This inadequate choice of odor stimuli is due to the dearth of affordable odor delivery systems that can generate plume-like, naturalistic stimuli with high reproducibility such as to allow for repeat measurements under laboratory conditions. We thus developed an odor delivery system that generates arbitrary time-varying patterns of individual odors and ternary mixtures at time scales of ~20 Hz. Here, we provide a detailed description of the construction and output characterization of our odor delivery system.

## 1. Introduction

The spatial and temporal dynamics of odors are hard to control given the volatile nature of odor stimuli. Also, odors are sticky and adhere to surfaces: this makes it difficult to deliver multiple odors in quick succession without any spillover between stimuli. Tight stimulus control is the first prerequisite for careful characterization of individual neuronal responses or animal behavior as a function of odor identity and concentration. Our goal was to design a low cost system that can deliver a wide range of time-varying odor stimuli with temporal statistics resembling natural stimulus dynamics (2-20 Hz) (Vickers 2000). At the same time, we aimed for the odor machine to reliably reproduce a given stimulus pattern multiple times to build a good estimate of the ‘average’ neuronal and/or behavioral response to a given stimulus pattern. Further, we required there to be no cross-contamination (between different odors or concentrations) when multiple stimuli are presented simultaneously or across consecutive trials. Finally, we optimized the machine for high stability in the output flow rate such as to avoid mechano-sensory stimulation of ORNs (Grosmaitre et al. 2007). Keeping this in mind, we designed our odor delivery system such as to fulfill four essential criteria:

*Criterion 1: Fast output kinetics to vary the output odor concentration at fast time scales (20 Hz).* A sharp, square pulse of odor is impossible to achieve given the slow diffusion rate of odors. Even if odor flow is gated through a fast-opening (millisecond kinetics) solenoid valve, odor output to the animal is slow (>tens of milliseconds). The output kinetics is governed by the diffusion rate and the velocity of the carrier stream. Thus to ensure fast kinetics, the odor of interest must be streamed through a high velocity carrier. However, gating a high flow rate air stream through any mechanical aperture (such as solenoid valves) imposes constraints on the stability of the output flow rate (*see **Criterion 4***).

*Criterion 2: High reproducibility of odor output.* Given the high variability in neuronal spiking and animal behavior, individual trials do not provide a reliable estimate of the neuron’s average response to a stimulus. To obtain an accurate estimate, the neuronal/behavioral response to the same stimulus must be measured a few times, while minimizing variability in stimulus delivery across repeated presentations. For odors, this translates into maintaining a stable, steady state output concentration despite different volatility of different odors, for the duration of the experiment (>1 hour).

*Criterion 3: Flexible and independent control of multiple odors with minimal cross-contamination between stimuli.* A frequent aim of olfactory studies is to understand how animals integrate inputs of different odors, arriving simultaneously (as a mixture) or sequentially. Such analysis requires independent control of multiple odors, without any cross-contamination between the presented odors. One way to ensure this is to route each odor through independent sets of tubing, valves etc such that a minimum number of components are shared between odors. However, this strategy becomes quickly unfeasible with increasing number of odors, given space and cost constraints. We therefore devised a lower-cost solution that enables fully independent delivery of up to three odors within a given experiment, while allowing flexible selection of any 3 odors (out of 20) across experiments.

*Criterion 4: Independent modulation of odor concentration without changing the output flow.* Olfactory receptors are mechano-sensitive. It is essential to alleviate mechanosensory cues to isolate only the chemo-sensory aspect of olfactory responses. Gating the odor output with a single solenoid valve inevitably changes the output flow rate. This is typically balanced by gating a flow-rate matched clean air stream through another solenoid valve, which opens and closes complementary to the odor valve. While this strategy stabilizes the net flow, it still results in relatively large instantaneous pressure transients at the switching time of the valves, particularly if odors are routed through a high velocity carrier stream (*as suggested in **Criterion 1***). Reducing the flow rate of the odor stream entering the valves significantly decreases these transients, but at the expense of slow output kinetics. The main design challenge for our delivery system was thus to achieve stable output flow without compromising output kinetics.

Keeping the above criteria in mind, we designed an odor delivery system that allowed us to present arbitrary time-varying odor patterns of individual odors (from a panel of 20) at a maximum frequency of 20 Hz. Here we describe in detail the construction and validation of this odor delivery system. We further discuss two adaptations of the single odor design that enable interleaved presentation of either multiple odors, or multiple concentrations of a given odor, with no spillover between odors/concentrations.

## 2. Methods

### Core odor machine design for delivering fast, time– varying stimuli of a single odor

1. *Creation of a saturated odorized stream of a select odor:* Saturated odor vapor was produced by bubbling the carrier air stream through undiluted, liquid odor contained in a custom-designed glass bubbler (Vensil, India) fitted with teflon corks (Figure 1a, *Odor vial*). Glass beads (VWR, 26396-506) were immersed in the odor to aid saturation and prevent aerosol formation. To achieve linear control of the desired output odor concentration, the odor-saturated carrier stream was serially diluted with charcoal filtered, humidified clean air at two consecutive stages in the odor machine – *Dilution 1* and *Final manifold*(Figure 1b). Odor selection was enabled through a digitally-controlled array of solenoid valves (Clippard ET-2M-12, **Figure S1a**) mounted on a linear manifold (Clippard, 15481-6) that channeled the carrier air stream through any one of six odor vials (Figure 1b, *Odor panel*). For a typical experiment, the carrier stream flow rate was regulated at 0.5 L/min using an acrylic block flow meter (Cole Parmer, EW-32460-40).
2. *10X dilution of the odor-saturated carrier:* The odor-saturated carrier stream was diluted 10X by mixing with a 5 L/min air stream (Cole Parmer, EW-32461-54, Figure 1b, *Dilution 1*). One-way check valves (Clippard MCV-1, MCV-1AB) were used to prevent back flow of the higher flow rate dilution air stream into the odor vial. A small fraction (0.5 L/min) of the 10X diluted odor stream was directed to the final valve assembly controlling odor delivery at the animal’s snout, and the rest was flushed to exhaust (Figure 1b, *Final manifold*).
3. *Second 10X dilution and final odor output:* Within the *final manifold*, the 10X diluted odor stream was directed towards the animal via manifold-mount solenoid valves (Clippard, ET-2M-12). The manifold was custom-designed such that the valve outputs were ejected into a continuous, high flow rate stream of clean air (5 L/min) which provided a second 10X dilution, bringing the final output concentration to ~1% saturation. To precisely control odor timing without altering the output flow rate, odor delivery was controlled by two pairs of solenoid valves *odor valves*(Figure 1b, *red*) and *air valves*(Figure 1b, *blue*). *Odor valves* shuttled the 10X diluted odor stream between the ‘Animal’ channel and ‘Exhaust’ channel. *Air valves* shuttled a flow-rate matched clean air stream between the ‘Exhaust’ and ‘Animal’ channel. The *odor* and *air* valves were turned ON-OFF in a complementary fashion (**Figure S1b**) such as to to keep the net output flow rate constant: balancing air was switched to the ‘Animal’ when odor went to ‘Exhaust’ (and vice-versa) (Figure 1b, *Final manifold*). To avoid pressure transients when the *odor* and *air* valves switched, it was critical to maintain a low flow rate (<0.5 L/min) for the odor/air stream shuttled by the solenoid valves. However, as discussed earlier, the low flow rate offered very slow kinetics, which was incompatible with our goal of delivering odor stimuli that fluctuate at ~20 Hz. To regain the fast kinetics at the animal’s snout, the ‘Animal’ channel was flushed with a steady stream of 5 L/min of clean air. Apart from accelerating output kinetics, this also provided a second dilution step bringing down the final output odor concentration to ~1% saturation. To maintain a constant back pressure in the whole system, the ‘Exhaust’ channel was also flushed continuously with a 5 L/min air stream. To further avoid pressure build-up, Valves within each pair (*odor* or *air*) were anti-coupled in hardware (**Figure S1b**), such the odor stream was kept continuously flowing throughout the experiment and simply switched from Animal→ Exhaust during an odor ON periods and from Exhaust→Animal during odor OFF periods. The continuous flushing of odorized air to exhaust significantly increases the rate of odor consumption per experiment (discussed later, Figure 3a). However, it offers two critical benefits: **a)** The continuous flushing of odorized air to exhaust during the stimulus period allows for clean (no pressure transients) and fast odor ON-OFF transitions. **b)** The continuous flushing of odorized air exhaust outside the stimulus period helps maintain a stable trial-to-trial output concentration. Note that the latter can in principle be substituted with long build-up periods preceding each stimulus, allowing the user to use more than one odor from the panel across different trials. However, in our experience, the optimal build-up duration is often longer than 10 seconds (depending on tubing length, odor used, etc) significantly increasing the inter-trial duration and consequently overall experiment duration. More importantly, the increase in total experiment time is further exacerbated by the need for additional ‘cleaning periods’ between consecutive trials to prevent cross-contamination between trials of two different odors. In the next section, we describe a modification of the core design that allows multi-odor experiments with no cross-contamination or penalization on total experiment duration.

**Figure 1:**
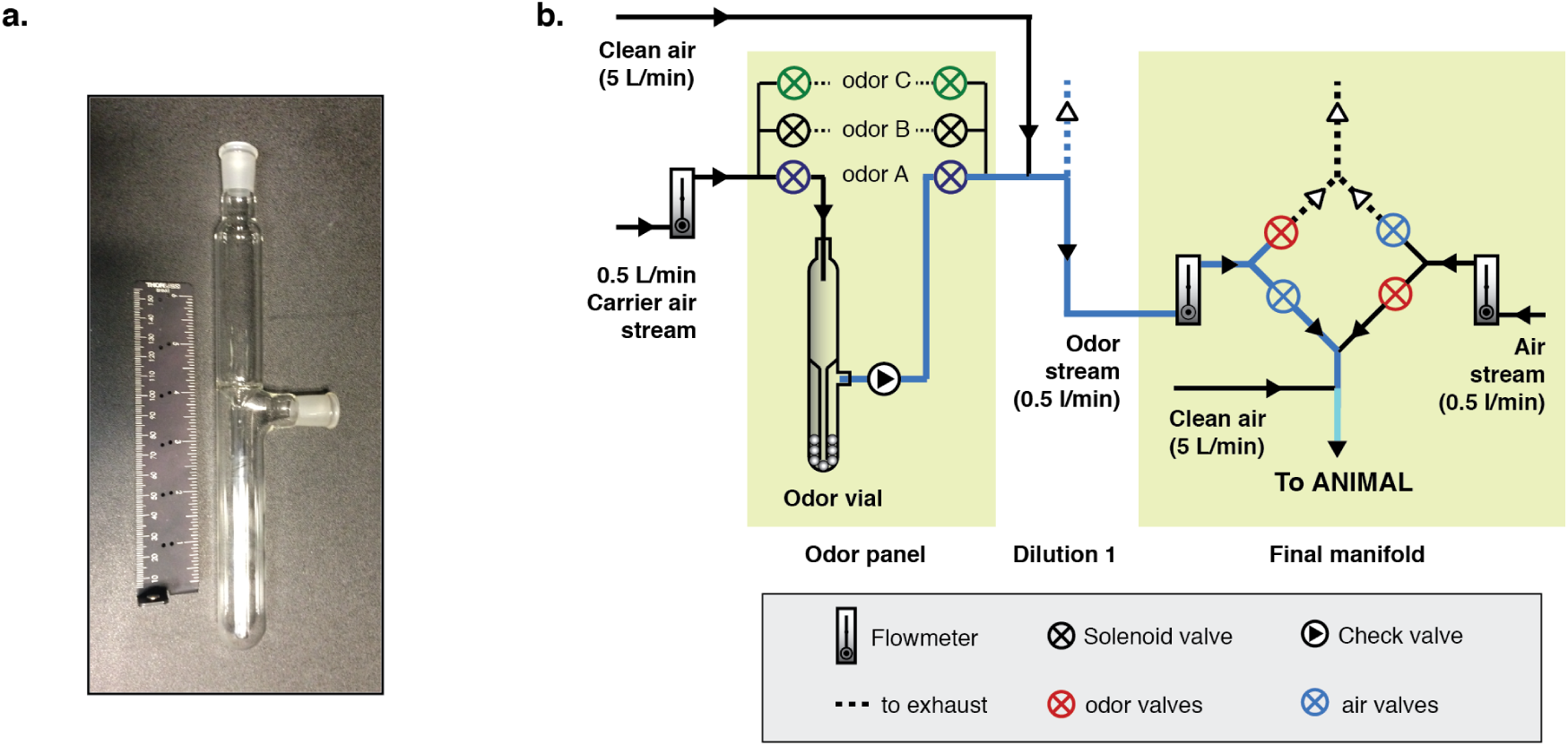
Schematic of odor delivery system for individual odors. **a:** Photograph of the glass bubbler used for holding undiluted odor. **b:** Schematic of the odor delivery system (*adapted from (Gupta et al. 2015)*). Saturated odor stream, produced by bubbling the carrier air stream (0.5 L/min) through a selected vial (e.g. Odor A) in the odor panel, is diluted 10 fold and directed to the final manifold at a regulated flow rate (0.5 L/min). Two pairs of anti-coupled solenoid valves allow rapid switching of the ‘odor’ and ‘flow rate matched clean air’ streams between Animal and Exhaust. A final 10 fold dilution by a fast carrier stream (5 L/min) ensures rapid kinetics and constant output flow. Pairs of valves that turned ON/OFF simultaneously are indicated in the same color.

The two-step serial dilution design reduces pressure transients and enables fast switching: Low flow rate of the odor stream at the switching point minimizes pressure transients as the odor valve is turned ON-OFF. The subsequent dilution with a high flow rate air stream allows for rapid odor clearance and fast stimulus kinetics. To further enhance the kinetics, we used short length (10 cm), narrow diameter tubing (4 mm) at the exit port, kept at a small distance (~1 cm) from the animal’s snout.

#### Modifications in system design for independent, simultaneous delivery of multiple odors

To deliver two or more odors simultaneously, with distinct temporal patterns, we added a minor modification to the core design. We used three independently controllable carrier streams, each serving an independent set (bank) of 6-8 odors (Figure 2). One odor was selected from each bank and diluted independently before routing to the final valve assembly at the animal’s snout. An independent balancing air stream was used for each odor. The final manifold was modified to accommodate 8 more pairs of valves, to follow the same switching system as described earlier for the single odor. The final valves were mounted on a custom-designed PEEK (polyetheretherketone) manifold (**Figure S2**, **Supplement – Manifold CAD file**) to minimize the dead volume, which allows for faster odor clearance time between trials.

**Figure 2:**
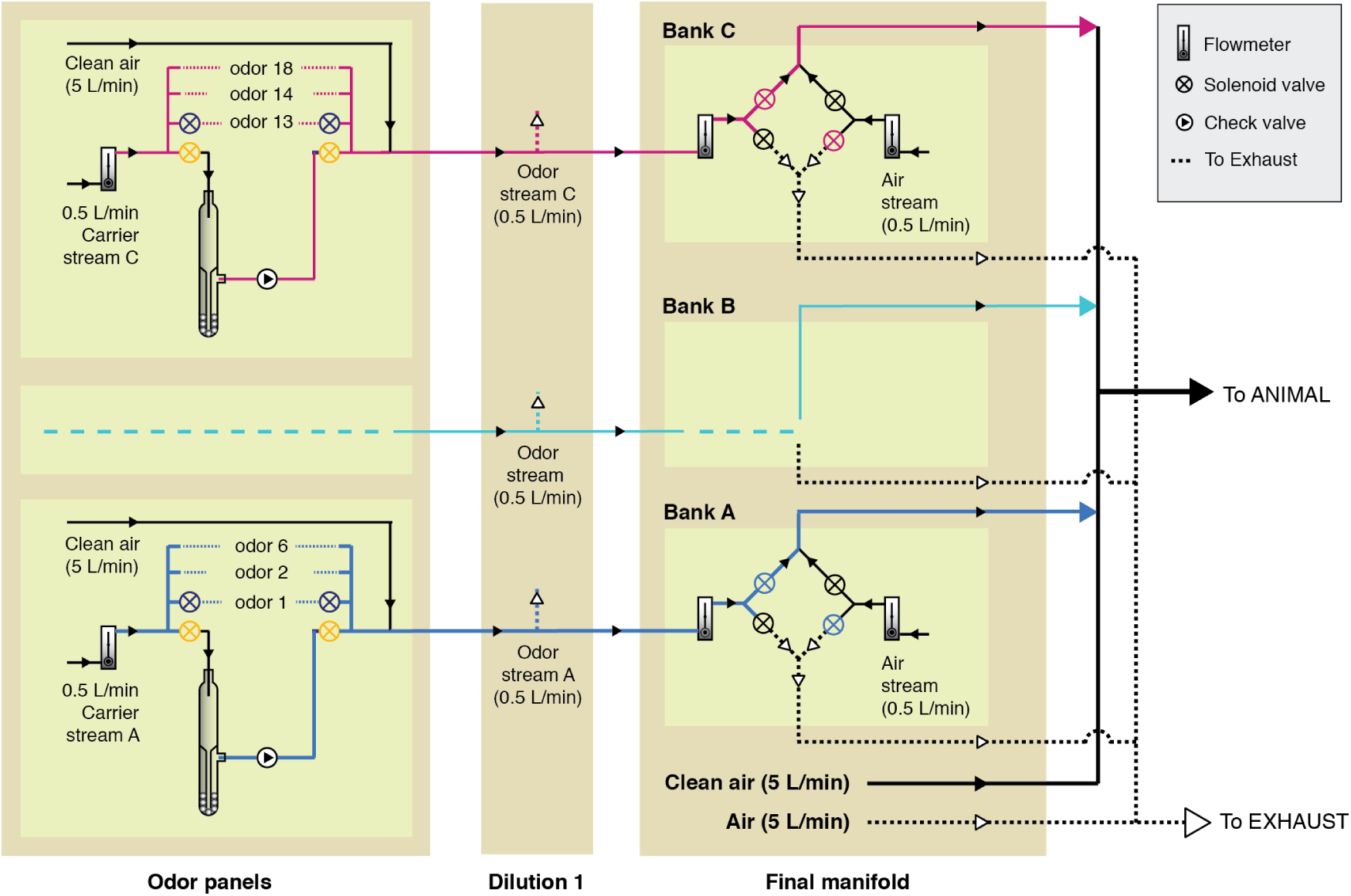
Schematic of odor delivery system for multiple odors and mixtures. Saturated streams of up to three different odors were produced by bubbling three independent carrier streams (0.5 L/min) through three distinct vials (one vial chosen per odor panel). The odor saturated streams were routed to independent dilution banks A, B, C (Dilution 1) for 10-fold dilution and further directed to the final valve assembly at the animal’s snout. In the final valve assembly, each odor stream was regulated by dedicated set of 4 valves (Bank A, B, C), which enable rapid and independent switching of the ‘odor’ and ‘flow rate matched clean air’ streams between ‘Animal’ and ‘Exhaust’. This scheme allowed us to deliver the three odors separately (one bank ON, other two flushed to exhaust), or as mixtures (two or more banks ON simultaneously).

#### Characterization of output odor concentration and flow rate

We used a Photo-ionization detector (PID, Aurora scientific, 200B miniPID) to characterize output odor concentration as a function of the odor valve ON-OFF state (Vetter et al. 2006). The output flow rate was measured simultaneously using an anemometer (Kurz instruments, 490-IS) inserted in the path of the output flow, ~2 cm from the outlet.

## 3. Results

### Reproducible odor output and no pressure transients

The PID response varied in synchrony with odor valve opening and closing. However, the output concentration dropped steadily from early trials to late trials as we presented several, randomly interleaved stimulus patterns mimicking a typical experimental session (12 trials per stimulus pattern spread over ~2 hours). The rate of decay differed from odor to odor (Figure 3a, *left*). We found that the concentration decay was also evident in reduced volume of odor remaining in the vial. Timely refilling of odor in each vial such as to maintain a fixed volume (8 ml) drastically reduced the inter-trial variability in the PID output (Figure 3a, *right*). With this simple strategy, the PID response was found to be highly reliable for a given odor and stimulus pattern, with low trial-to-trial variability (across 1-2 hours of constant usage of the odor machine) even when trials of many stimulus patterns were interleaved (Figure 3a, Figure 3b). In contrast, the anemometer output was unaffected by valve opening and closing, indicating a stable output flow rate (Figure 3b). For any given odor in our panel, we were able to predict the odor output for any arbitrary stimulus pattern by assuming a generic PID response function and convolving it with the time-series of odor valve opening and closing (**Figure S3a**).

**Figure 3:**
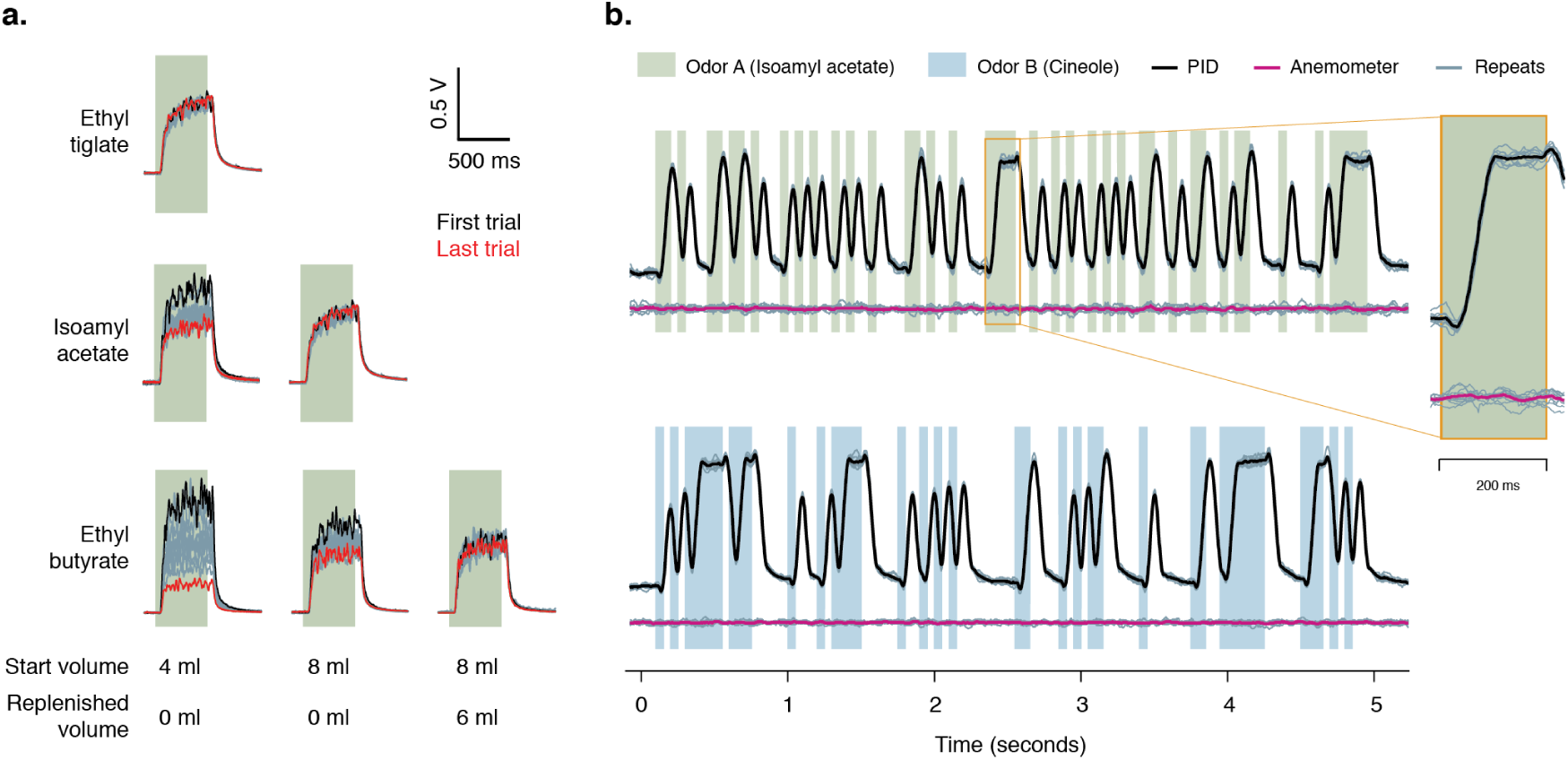
Reproducible and predictable odor output for arbitrary time-varying odor stimulus patterns. **a:** Comparison of output reproducibility for 500 ms long pulses of three odors, across 12 trials, randomly interspersed in a long experiment session (~2 hours). Grey lines show all 12 trials. Black and red lines show the first and the last trial. Concentration drops across 2 hours more steeply for Ethyl butyrate than Ethyl tiglate. Using larger volumes of initial odor load (8ml versus 4 ml) and mid-experiment replenishment reduces output variability. **b:** Observed output profile for Isoamyl acetate (*reproduced from (Gupta et al. 2015)*) and 1,4-Cineole (1% saturation) for pseudo-random sequence of odor pulses. Vertical green and blue bars mark odor valve ON periods. Black and Pink lines show simultaneously measured, average read-outs of a photo-ionization detector (PID, black) and Anemometer (Pink) (sampling rate 1 KHz). Sensor outputs were measured in volts and are plotted here in arbitrary units (a.u.). Grey lines show individual trials (10 trials). Inset shows enlarged view of a 200 ms long pulse of Isoamyl acetate within the sequence.

### Fast kinetics for odor build-up and clearance

The average ON and OFF kinetics for the odors in our panel were found to be 40.1*±*2.8 ms and 71.9*±*8.7 ms respectively, and were similar across odors (Figure 4). The dead time was also consistent across odors (31.6*±*1.4 ms) (Figure 4b). As discussed earlier, the output kinetics increased with higher net output flow rate (**Figure S3b**).

**Figure 4:**
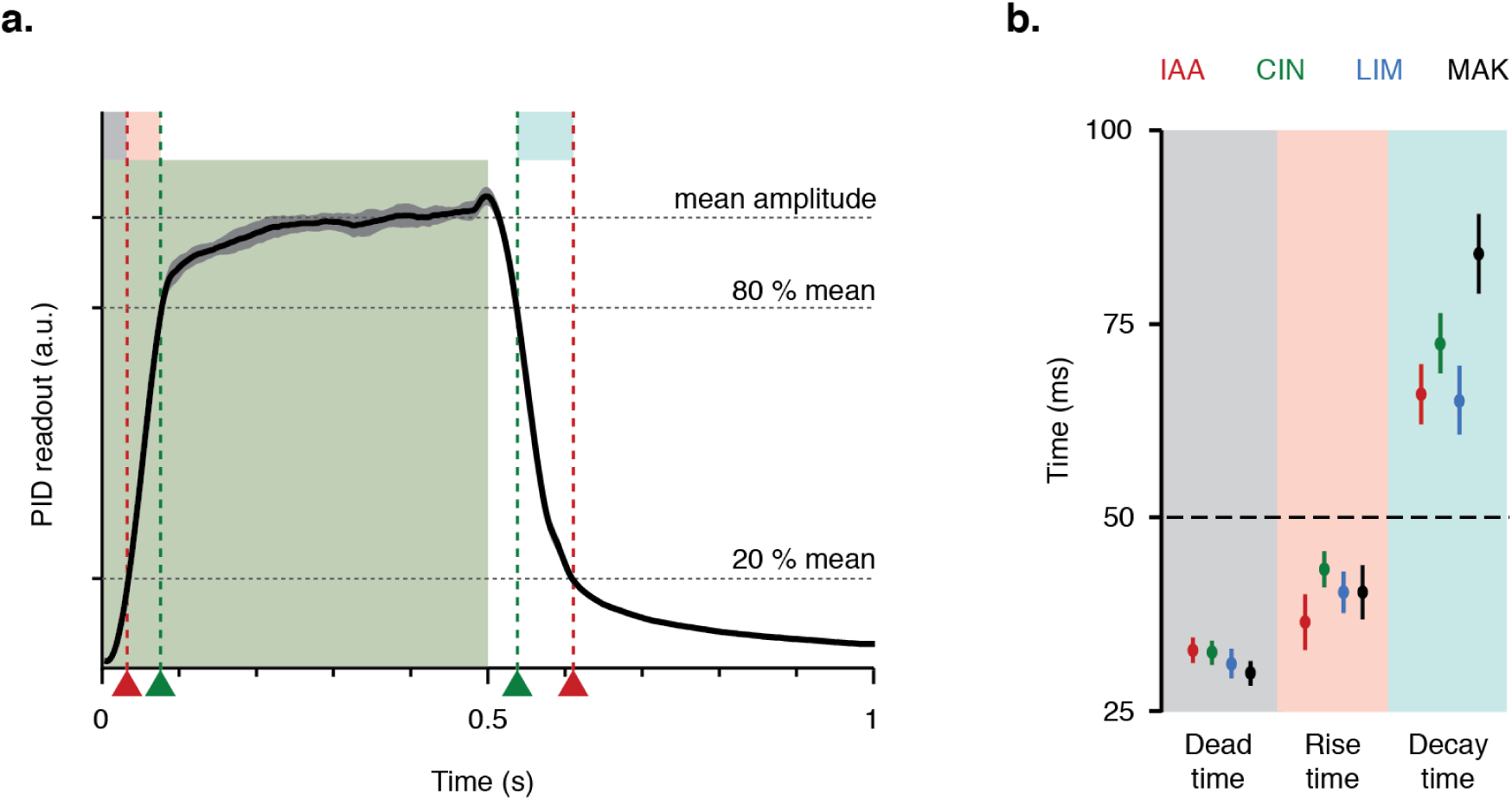
Fast output kinetics of the odor delivery system across chemically diverse odors. *Figure reproduced from (Gupta et al. 2015)* **a:** Photo-ionization detector (PID) output profile for a 500 ms pulse of Isoamyl Acetate (1% saturation). Vertical green bar marks odor ON period. Black line shows average response (12 trials, sampling rate 320 Hz). Grey band shows one standard deviation. Red and green lines indicate time taken to reach 20% and 80% of the mean odor amplitude respectively. Dead time (grey) is the time taken from valve opening to reach 20% of mean. Rise time (pink) and decay time (blue) are the time taken to reach from 20% to 80% of mean amplitude and vice versa respectively. **b:** Dead time, rise time and decay time for Isoamyl Acetate (IAA), Cineole (CIN), Limonene (LIM) and Methyl Amyl Ketone (MAK) at 1% saturation. Error bars indicate one standard deviation (pulse duration 500 ms, 20 repeats).

### No cross-contamination across odors for binary odor stimuli

We interleaved presentations of stimulus patterns from two odors individually as well as their binary combinations. We found that the PID amplitude during periods when both odors were presented could be reliably predicted as the sum of PID responses evoked by each odor presented individually, irrespective of the temporal pattern for each stimulus (Figure 5). This confirmed that the output for two odors was indeed independent and mixtures were a linear sum of the composing odor stimuli.

**Figure 5:**
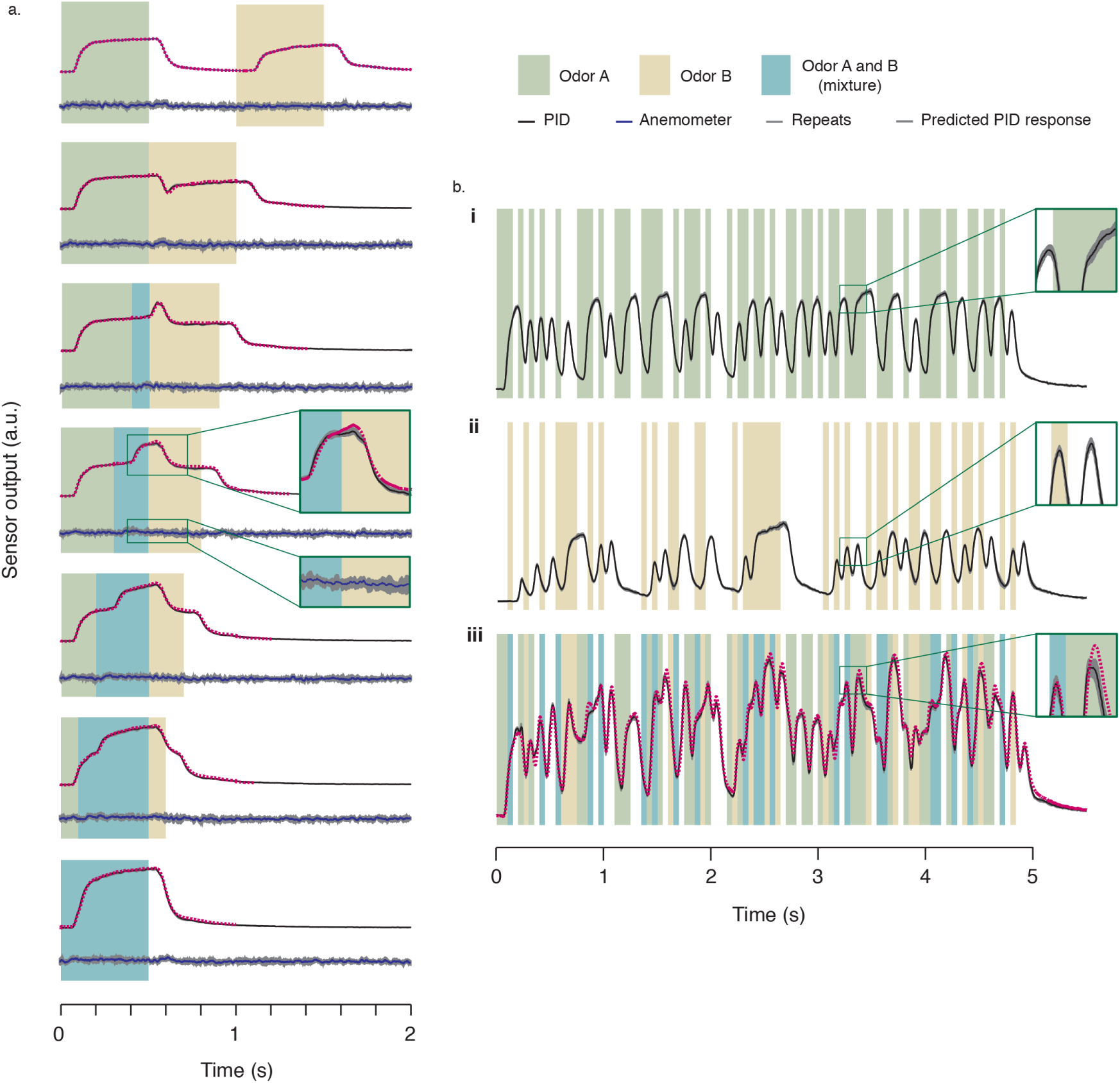
Output odor characteristics for binary odor stimuli. *Figure reproduced from (Gupta et al. 2015)* **a:** PID and anemometer output profile for pairs of odor pulses of Isoamyl acetate (IAA, 1% saturation) and Limonene (LIM, 1% saturation) at varying inter-pulse intervals. Vertical green, yellow and cyan bars represent odor ON periods for IAA, LIM, or both, respectively. Black and blue lines show simultaneously measured, average PID and anemometer response respectively. Grey bands indicate one standard deviation (10 trials). Dotted red lines show the expected PID output, calculated as a sum of the measured PID outputs for individual pulses of each odor. Inter-pulse intervals from top to bottom (in ms): 1,000 (no overlap), 500, 400, 300, 200, 100 and zero (complete overlap); individual pulse durations: 500 ms. **b:** PID output profile for pseudo random fluctuating patterns of two odors presented simultaneously. **b(i,ii).** Black lines show average PID output for a fluctuating pattern of Limonene (LIM, 1% saturation) and Cineole (CIN, 1% saturation) respectively. **b(iii).** Black and dotted red lines show observed and expected PID response upon simultaneous presentation of the patterns in **b(i)**and **b(ii)**. Grey bands indicate one standard deviation (10 trials each). Vertical green, yellow and cyan bars represent odor ON periods for LIM, CIN, or both, respectively.

### Odor delivery system for linear and inter-leaved concentration control

Olfactory studies frequently require precise control of odor concentration on a trial-to-trial basis. While individual experimental demands may vary, we outlined three broad criteria that must be met by an ideal odor delivery system for concentration control. Below, we describe these criteria and the design solutions used in our odor delivery system that enable us meet these criteria.

*Criterion a: Obtain a desired concentration series (e.g. 1X, 5X, 10X) irrespective of the chemical nature of the selected odor.* Typically, increasing odor concentrations are obtained by flushing increasing flow rates through the odor vial via a mass flow controller (MFC). We found that the range of concentrations produced by this approach was highly dependent on odor chemistry, and often resulted in unexpectedly supra-linear odor output at lower flow rates for viscous odors (**Figure S4**). We bypassed this limitation by using only one saturation step (flow through the odor vial) for each odor and distributing the saturated odor stream into different concentration banks (*Dilution1*, *Dilution2* etc.) via multiple serial dilutions (Figure 6a). As a result, the relative ratio between the outputs of different concentration banks was purely dictated by the dilution factor and did not depend on individual odor chemistry (Figure 6b-c).

**Figure 6:**
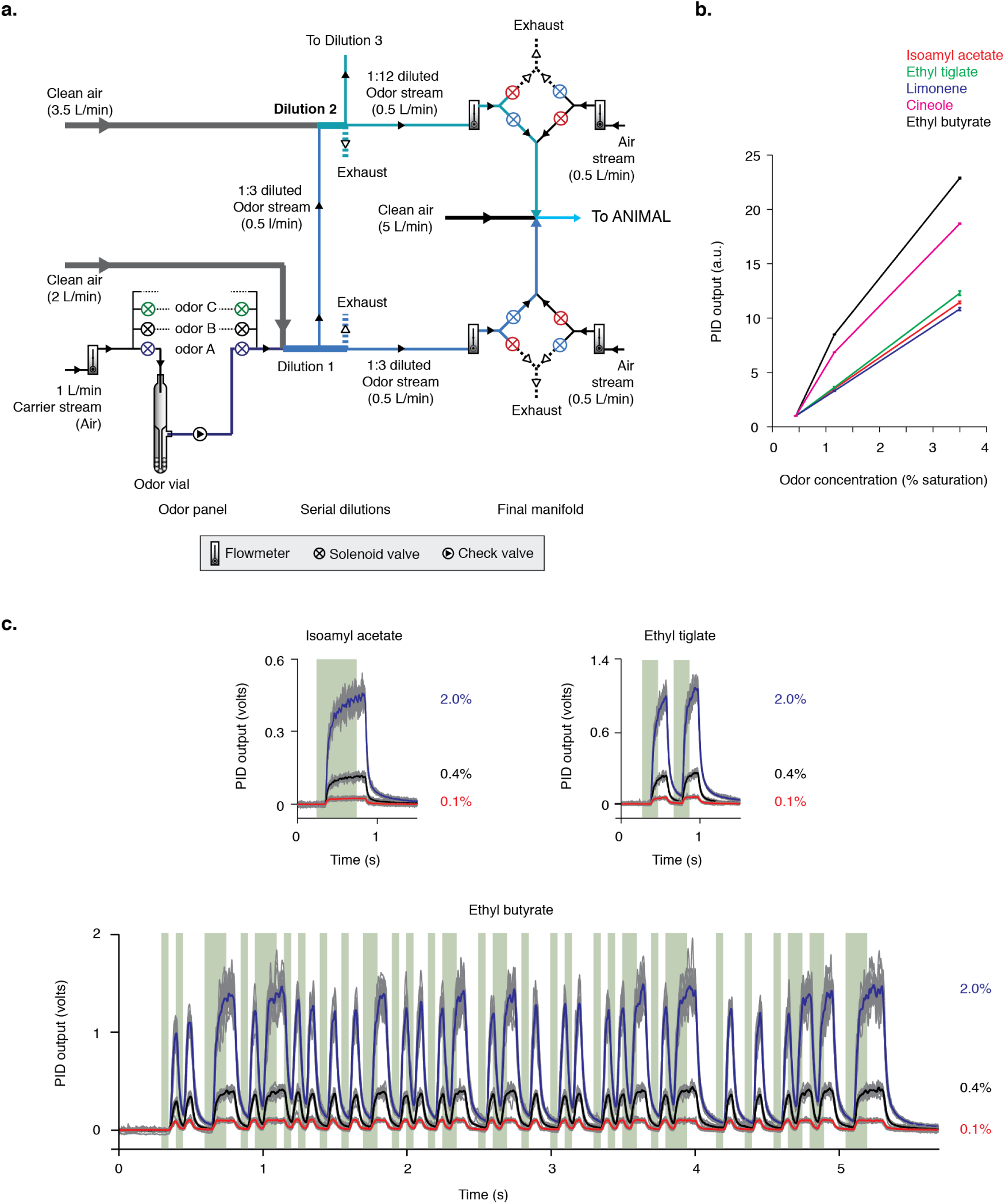
Linear, inter-leaved concentration control across chemically diverse odors. *Figure reproduced from (Gupta et al. 2015)* **a:**Schematic of the odor delivery system for reliable and linear concentration control. Saturated odor stream, produced by bubbling the carrier air stream through a selected vial (e.g. Odor A) in the Odor panel is diluted with a 2 L/min clean air stream to obtain 1:3 dilution. One fraction of this 1:3 diluted odor stream is routed to the final manifold at a regulated flow rate (0.5 L/min) where it is further diluted 10-fold by a high-flow rate carrier stream (5 L/min) and switched between Rat and Exhaust by two pairs of anti-coupled solenoid valves (similar to that described in Figure 1b). This results in a final output concentration of 3.5% saturation at the animal’s snout. Lower concentrations of the same odor are obtained by setting up additional serial dilutions of the initial 1:3 diluted odor stream before the final manifold. For example, a second fraction of the 1:3 diluted stream is mixed with 3.5 L/min clean air to obtain a net dilution of 1:12 instead of the original 1:3 dilution. This 1:12 dilution stream is also routed to the final manifold at a regulated flow rate (0.5 L/min) and switched between ‘Animal’ and ‘Exhaust’ by the same mechanism as that described for the 1:3 diluted stream. As a result, the net output concentration of this stream is 4 times lower that the first stream. Even lower concentrations can be obtained by setting up as many serial dilutions as required, of the original 1:3 diluted stream. **b:**) Linear odor output across five chemically diverse odors measured as the average photo-ionization detector (PID) response amplitude within a 500 ms odor pulse. Average PID amplitude was calculated from 12 trials across randomly interleaved presentations of three different concentrations. Error bars indicate one standard deviation of the mean. **c:** Observed output profile for three odors (Isoamyl acetate, Ethyl tiglate and Ethyl butyrate) for stimulus patterns delivered at three different concentrations (0.1%, 0.4% and 2% saturation). Vertical green bars mark odor valve ON periods. Red, black and blue lines show average response amplitude of a PID (sampling rate 1 KHz) across 12 trials at three different concentrations (0.1%, 0.4% and 2%) from a set of randomly interleaved trials of all three concentrations. Grey lines show individual trials. Note that the relative difference in amplitude across the three concentrations for each odor is similar despite the differences in PID sensitivity for each odor.

*Criterion b: Interleave presentations of different concentrations in quick succession, with no cross-contamination between concentrations.* A common challenge faced when trying to interleave multiple odor concentrations is significant spill-over of residual odor from a preceding high odor concentration trial to the odor current trial, raising the output concentration of what may be intended as a ‘low’ odor concentration trial. To avoid spill-over between desired odor concentrations, we turned to the same solution as described earlier for delivery of multiple odors with no cross-contamination. We dedicated one odor bank to each desired odor concentration and treated the output of each concentration bank as an independent odor, using a dedicated set of solenoid valves for each bank (Figure 6a). Thus, a low concentration trial could be presented immediately following a high concentration trial with no spillover of odor from the previous trial. This can be appreciated in the low inter-trial variability of the observed PID responses for different concentration trials, even when all concentrations were presented in a randomly interleaved sequence (Figure 6c).

*Criterion c: Low variability in the absolute odor concentration.* As discussed earlier, we found that depending on the volatility of different odors, the concentration may drop to half its original amplitude within 30 minutes of continuous usage (e.g. Ethyl butyrate). However, replenishment of the odor volume to match the initial amount in the vial immediately restored the PID readings to their original values. We therefore independently characterized the decay in odor concentration for each odor, and replenished odor volumes in the vial at the appropriate frequency for each odor (Ethyl butyrate @ every 15 minutes, Ethyl tiglate @ every 30 minutes). This can also be appreciated in the low variability across individual trials in the example PID responses in Figure 6c. The individual trials (grey lines) for each concentration are randomly interspersed across duration of 1.5-2 hours. Additionally, for the higher concentrations, care was taken to replace the valves periodically as the high concentration odors frequently clogged both the solenoid and check valves. For the same reason, we restricted the highest concentration tested in our experiments to 3.5%. For the odors we sampled, at concentrations >3.5%, valves got clogged even within one experimental session.

In summary, we developed an odor delivery system that can deliver rapidly fluctuating stimulus patterns of individual odors and their mixtures with high reproducibility across hundreds of trials. Given the fast output kinetics (40.1*±*2.8 ms), it is ideal for generating naturalistic plume-like odor stimuli in controlled laboratory settings. It also allows sequential presentation of multiple odor concentrations with high predictability and no cross-contamination between trials of high and low concentrations.

## 4. Discussion

Difficulties in precise control of olfactory stimuli have been a long-standing challenge in olfactory research(Vickers et al. 2001; Vetter et al. 2006). While recent advances in optogenetic strategies combined with patterned illumination (Dhawale et al. 2010) offer one approach to overcome some of these limitations, they do not replicate natural stimulus dynamics and are limited to selective stimulation only on the dorsal bulb surface. The tight control of stimulus conditions is critical to accurately estimate the relationship between features of odor stimuli and neuronal responses (Gupta et al. 2015) and/or animal behavior. These conditions include the lack of flow transients during odor ON-OFF transitions, reproducibility of odor amplitudes and time-courses, and rapid kinetics to prevent odor spillover between pulses delivered in close succession.

A few elegant odor delivery systems have been described recently for precise control of odor concentration of individual odors (Kim et al. 2011; Martelli et al. 2013). The odor delivery system described in this chapter differs from the previously available systems in a three key aspects: first, faster output kinetics (~20 Hz) while maintaining a stable output flow rate; second, inter-leaved presentation of different odors or different concentrations of a given odor, even within a single trial, with negligible cross-contamination; and third, linear control of output concentration across chemically diverse odors. This was made possible by use of a multi-step serial dilution design, which exploits a combination of low input flow rates to reduce output pressure transients and high output flow rates to achieve fast output kinetics. The linearity of the concentration output (Figure 6b) is conferred by the use of a common step for creating 100% saturated vapor of the odor and modulating concentration only via serial dilutions of the odorized air. The ability to interleave different odors/concentrations is conferred by the fully independent control of each odor/concentration stream at the final manifold. Since the different odor/concentrations do not share any common valves, there is no cross-contamination between odors/concentrations. As a result, we were able to deliver low concentration odor stimuli (0.5% saturation) in quick succession to high concentration (3.5% saturation) ones without spillover across trials.

A major limitation of the odor delivery system described in this study is the small number of odors/concentrations (maximum 3) that can be interleaved within the same experiment (consecutive trials)-one odor/concentration per bank (Note that across experiments, different odors/concentrations can be used on the same bank by adequate flushing of residual odor). Similar limitation applies to the currently described design for concentration control. While the overall stimulus panel is large (several odors in the odor panel), multiple concentrations of only one user-selected odor can be interleaved within a given experiment. However, the odor machine design is modular and can be extended to deliver more than three odors/concentrations by addition of more banks. If experiment duration is not limiting, another possible strategy to increase odorant/concentration diversity is to switch between multiple odors on the same bank between consecutive trials. However, care must be taken to allow a sufficiently long inter-trial period such as to fully flush out residual odor from the previous trial. Routing a high flow rate air stream (cleaning line) can significantly aid this process. Note however that some residual odor will still remain in the final odor valve that directs the diluted odor stream on each bank towards the animal. This residual odor can only be flushed by routing clean air instead of odor through the valve. By design, this will result in delivery of the residual odor to the animal outside the designated odor period – a condition that is typically incompatible with behavioral experiments. Thus, the ideal solution is to set up independent banks for each odor-concentration pair in the stimulus set. While each additional bank does increase system complexity and cost, the use of manual flow meters instead of MFCs in our design, significantly reduces the overall cost and makes the design expansion more affordable.

In summary, the odor delivery system described here offers a low-cost, integrated solution for interleaved presentations of deterministic, arbitrary time-waveforms of individual odors, mixtures and concentrations at fast time scales (20 Hz). It is ideally suited for behavioral experiments that require inter-leaved presentations of multiple odors (or concentrations) and rely on careful concentration control, while mimicking natural stimulus dynamics.

## 5. Additional Methods

### Odors Used

We tested a total of 9 odors - Isoamyl acetate (W205532, Sigma-Aldrich), 1,4 - Cineole (W365807, Sigma-Aldrich), Limonene (W504505, Sigma-Aldrich), Methyl amyl ketone (W254401, Sigma-Aldrich), Amyl acetate (W504009, Sigma-Aldrich), Ethyl tiglate (W246018, Sigma-Aldrich), *γ*-Terpinene (W355909, Sigma-Aldrich), Linalool (W263508, Sigma-Aldrich), Ethyl butyrate (W242705, Sigma-Aldrich). The odors were chosen on the basis of detectable Photo-ionization detector (PID) signals and rapid clearance from the solenoid valves and tubing, while maintaining a diverse range of functional groups in our odor stimulus panel. Several odors sampled, such as 1-Octanol, 1-Hexanol, Citral etc. did not evoke reproducible PID responses, and were excluded from the stimulus panel.

### Data analysis

Analysis was done in Matlab (Mathworks) with custom written routines.

### Figure 4

To characterize the output kinetics, we generated 500 ms long odor pulses and calculated the mean PID response amplitude in the later half of the valve ON period (20 repeats per odor). We defined the ON kinetics as the time taken to rise from 20% to 80% of the mean PID amplitude after valve opening. The OFF kinetics was conversely defined as time taken to fall from 80% to 20% of the mean amplitude post odor valve closing. The dead time was defined as the fixed delay to reach 20% mean odor amplitude after valve opening.

### Figure 6b

To characterize output linearity (Figure 6b) across multiple concentrations, we measured the mean odor amplitude for 500 ms long odor pulses (12 repeats) from randomly interleaved presentations of three different concentrations of each odor.

### Parts list

See Supplementary material – Table 1. See **Supplement** for a 3D CAD drawing of the custom designed manifold for the final valve assembly.

### Software for odor machine control

The odor delivery system was controlled via custom written LAB-VIEW code. The code is available at https://github.com/priyanka-cshl/odor_machine_control.git.

## 6. Acknowledgments

We would like to thank K. Parthasarathy and A. Dhawale for insight and help in designing the odor delivery system and Fred Marbach for comments on the manuscript and Latex tutoring. Dev Kumar (Machinist, NCBS) and Rob Eifert (Machinist, CSHL) machined the custom parts described in this study.

## 8. Supplement

Supplemental material

- Supplementary figures S1-4
- Table 1: Odor machine parts list

Manifold CAD file: final_valve_manifold.zip

